# Molecular detection of *Leishmania donovani, Leishmania major*, and *Trypanosoma* spp. in *Sergentomyia squamipleuris* sandflies from a visceral leishmaniasis focus in Merti sub-County, eastern Kenya

**DOI:** 10.1101/2020.07.29.226191

**Authors:** Barrack O. Owino, Jackline Milkah Mwangi, Steve Kiplagat, Hannah Njiriku Mwangi, Johnstone M. Ingonga, Alphine Chebet, Philip M. Ngumbi, Jaundouwe Villinger, Daniel K. Masiga, Damaris Matoke-Muhia

**Author notes:** Equal first authors (Barrack O. Owino, Jackline Milkah Mwangi).

## Abstract

**Background:** Visceral leishmaniasis (VL) and zoonotic cutaneous leishmaniasis (ZCL) are of public health concern in Merti sub-County, Kenya, but epidemiological data on transmission, vector abundance, distribution, and reservoir hosts remains limited. To better understand the disease and inform control measures to reduce transmission, we investigated the abundance and distribution of sandfly species responsible for *Leishmania* transmission in the sub-County, and their blood-meal hosts.

**Methods:** We conducted an entomological survey in five villages with reported cases of VL in Merti sub-County, Kenya, using CDC miniature light traps and castor oil sticky papers. Sandflies were dissected and identified to the species level using standard taxonomic keys and PCR analysis of the cytochrome c oxidase subunit 1 (COI) gene. *Leishmania* parasites were detected and identified by PCR and sequencing of internal transcribed spacer 1 (ITS1) genes. Bloodmeal sources of engorged females were identified by high-resolution melting analysis of vertebrate cytochrome b (cyt-b) gene PCR products.

**Results:** We sampled 526 sandflies consisting of eight species, *Phlebotomus orientalis* (1.52%; n = 8) and seven *Sergentomyia* spp. *Sergentomyia squamipleuris* was the most abundant sandfly species (78.71%; n = 414) followed by *Sergentomyia clydei* (10.46%; n = 55). *Leishmania major, Leishmania donovani*, and *Trypanosoma* DNA were detected in *S. squamipleuris* specimens. Humans were the main sources of sandfly bloodmeals. However, we also detected mixed bloodmeals; one *S. squamipleuris* specimen had fed on both human and mouse (*Mus musculus*) blood, while two *Ph. orientalis* specimens fed on human, hyrax (*Procavia capensis*), and mouse (*Mus musculus*) blood.

**Conclusions:** Our findings implicate the potential involvement of *S. squamipleuris* in the transmission of *Leishmania* and question the dogma that human leishmaniases in the Old World are exclusively transmitted by sandflies of the *Phlebotomus* genus. The presence of *Trypanosoma* spp. may indicate mechanical transmission, whose efficiency should be investigated. Host preference analysis revealed the possibility of zoonotic transmission of leishmaniasis and other pathogens in the sub-County. *Leishmania major* causes ZCL while *L. donovani* is responsible for VL. However, the reservoir status of the parasites is not uniform. Further studies are needed to determine the reservoir hosts of *Leishmania* spp. in the area.

## Introduction

Cutaneous leishmaniasis (CL) and visceral leishmaniases (VL) are vector-borne parasitic diseases that are caused by protozoan parasites of the *Leishmania* genus and transmitted by infected female sandflies during a bloodmeal [1]. Cutaneous leishmaniasis and VL are among the world’s most neglected tropical diseases (NTDs), occurring mainly in remote foci of the tropical, subtropical, and Mediterranean regions. Approximately 350 million people are at risk of infection [2]. Cutaneous leishmaniasis is the most common form with over 600,000 annual cases worldwide [3] and contributes to high psychological morbidities due to its scarring and stigmatising lesions [4]. In contrast, VL is the most fatal form of leishmaniasis with a fatality rate of nearly 100%, which occurs within two years if left untreated [3].

In Kenya, both VL and CL are endemic in arid and semi-arid areas in the Rift Valley, eastern and north-eastern regions in the country [5]. Cutaneous leishmaniasis is caused mainly by three species of *Leishmania*: *Leishmania tropica, Leishmania major*, and *Leishmania aethiopica* [2,6,7]. Cutaneous leishmaniasis infections caused by *L. aethiopica* and *L. tropica* are predominant in highland areas, especially around the Mount Elgon and Rift Valley regions, respectively [7]. In contrast, *L. major* infections and VL, caused by *L. donovani*, are common in low altitude arid and semi-arid areas of the Rift Valley, eastern and north-eastern Kenya [5,8].

Sandflies of the *Phlebotomus* genus are the confirmed vectors of human leishmaniasis in the Old World [1]. In Kenya, the sandflies *Phlebotomus guggisbergi, Phlebotomus duboscqi*, and *Phlebotomus pedifer* are the confirmed vectors of *L. tropica, L. major*, and *L. aethiopica*, respectively [2,9]. In contrast, *L. donovani* is transmitted primarily by *Phlebotomus martini*, while *Phlebotomus orientalis* is regarded as a secondary vector of the parasite in Kenya [9,10]. Sandfly species belonging to the *Sergentomyia* genus are regarded as non-vectors of human leishmaniases since some species within this genus do not support the development of *Leishmania* parasites within their midgut [11,12]. However, *Sergentomyia* sandflies are the predominant species in most leishmaniasis endemic foci in the Old World, where they appear to be well adapted to different biotopes and environmental conditions [13]. Most of these species have been shown to feed on humans and other mammalian hosts [2] which may carry other pathogens in addition to *Leishmania* species.

Various studies have recently suggested the possible role of some *Sergentomyia* species in the transmission of *Leishmania* parasites pathogenic to humans. For instance, studies conducted in CL hotspot in other countries revealed the presence of *L. major* DNA in *Sergentomyia sintoni, Sergentomyia darlingi*, and *Sergentomyia minuta* [13–16]. Furthermore, Mukherjee and colleagues identified *L. donovani* infections in *Sergentomyia babu* in India [17]. Other reports from a CL outbreak area in Ghana showed that *L. major, L. tropica*, and *Trypanosoma* spp. can infect *Sergentomyia ingrami, Sergentomyia hamoni*, and *Sergentomyia africana africana* sandflies, respectively [18]. Live *L. major* promastigotes have also been isolated from *Sergentomyia garnhami* in Kenya [19].

Knowledge of vectors, feeding preference, and the *Leishmania* species in circulation are crucial for the control of leishmaniasis transmission in disease-endemic areas. However, these data are scarce for Merti sub-County where repeated outbreaks of leishmaniasis have been reported. We designed this study to identify the sandfly species with the potential to transmit *Leishmania* parasites in Merti sub-County, Kenya, as well as determining their abundance, distribution, and bloodmeal sources. The information would contribute to a better understanding of the disease and the implementation of targeted vector control measures.

## Materials and Methods

### Study area

We conducted an entomological survey in July 2018, in five villages in Merti sub-County (1.066488° N; 38.657372° E) (**Fig**.**1**). We selected this area based on recurrent leishmaniasis outbreaks and an upsurge in the number of suspected VL cases among the patients visiting local health facilities in Merti. The sub-County is situated in Isiolo County at an altitude of 347 metres above sea level and approximately 420 km to the north-east of Nairobi. It covers an area of approximately 12612 km^2^ which is entirely arid or semi-arid with a population of 47206 people according to the 2019 national census [20]. The sub-County has two rainfall seasons [21]. The long rains occur between March and May, while the short rains, which is the most significant, start from October to December. The annual rainfall in the area ranges between 100-250 mm with an average annual temperature of 29°C [21]. Livestock production is the main economic activity in the area with nomadic pastoralism being prominent.

### Sample size determination and sandfly sampling

Since sandflies for this study were sampled at a single time point in time (cross-sectional sampling), we estimated the sample size, using the prevalence formula, according to *Amin et*.*al* [23]. Due to the lack of information on *Leishmania* infection prevalence in the vectors, we calculated our sample size assuming an expected infection prevalence of 50% to obtain an optimum sample size for the study. Therefore, at 95% confidence level and 5% precision, we estimated that a minimum of 385 sandflies was required to satisfy the study objectives.

Sandflies were sampled from five villages in Merti sub-County (Barsa, Kambi Juu, Korbesa, Malkagala and Matt Arban) using 12 CDC miniature light traps (John W. Hock Co., Gainesville, FL, USA) and castor oil sticky paper traps. The traps were placed in animal sheds and outdoors in areas likely to harbour sandflies such as near the *Acacia* trees and in chromic vertisols where *Ph. orientalis* and other sandflies are thought to breed and/or rest [10]. None of the traps was set indoors. We set the traps at 0.5-1.5 metres above the ground in each of the sampling sites from 1800 hrs to 0600 hrs the following day to constitute one trapping night. This was repeated over a period of 12 days.

### Sandfly dissections and morphological species identification

Following each trapping night, sandflies were sorted and washed in 2% detergent followed by antibiotic and antifungal solutions [24]. For all the sandflies, the head and the third last abdominal segment were dissected and cleared in gum chloral hydrate, which also served as the mountant, for morphological species identifications [25]. The abdominal status of the female sandflies was recorded as fed, unfed or gravid. Furthermore, we examined the midguts of individual female sandflies for the presence of live promastigotes. Morphological species identification of the mounted sandfly specimens was based on the external genitalia of males and features of the pharynx, antennae and spermatheca for females using standard taxonomic keys [26–28]. The remaining parts of the dissected female sandflies (i.e. the thorax, wings, legs, and abdomen) were preserved in 70% ethanol and transported under dry ice to the Centre for Biotechnology Research and Development (CBRD), KEMRI for parasite culture and further morphological identification, and International Centre of Insect Physiology and Ecology (*icipe*) laboratories for molecular analyses.

### Molecular identification of sandfly species

In the laboratory, we homogenised the remaining parts of each sandfly specimen in 180 μl of buffer ATL (QIAGEN, Hannover, Germany) taking care to avoid contamination between the specimens. Genomic DNA was extracted from each homogenate using the DNeasy Blood and Tissue Kit (QIAGEN, Hannover, Germany) according to the manufacturer’s recommendations. The extracted DNA was stored at -20°C until use.

To validate the morphological species identifications, we amplified the sandfly mitochondrial cytochrome c oxidase subunit I (COI) gene according to Kumar *et. al* using the primers; forward LCO 1490 (5’-GGTCAACAA ATCATAAAGTATTGG-3’) and reverse, HCO 2198 (5’-TAAACTTCAGGGTGACCAAA AAATCA-3’) [29]. The resulting 700 bp *COI* amplicons were purified using the QIAquick PCR purification kit (QIAGEN, CA. USA) according to the manufacturer’s recommendations and submitted to the Macrogen (The Netherlands) for sequencing using the forward primer.

### Detection of *Leishmania* and *Trypanosoma* species in sandflies

We analysed 57 sandfly samples consisting of all the 40 blood-fed and 17 unfed samples for the presence of *Leishmania* DNA. The blood-fed samples consisted of *Sergentomyia squamipleuris* (90%, n = 36) and *Ph. orientalis* (10%, n = 4) while the representative unfed samples comprised of *S. squamipleuris* (82.3%, n = 14), *Ph. orientalis* (11.8%, n = 2) and *Sergentomyia clydei* (5.9%, n = 1).

We carried out the detections in individual specimens by PCR amplification of the *Leishmania* internal transcribed spacer 1 (ITS1) region using L5.8S and LITSR primers [30]. The 20-μl PCR mixtures contained 1x Dream Taq buffer with 2 mM MgCl_2_ (Thermo Scientific, USA), 0.25 mM dNTPs mix, 500 nM of each primer, 2 U of Dream Taq DNA polymerase (Thermo Scientific, USA), 5–10 ng of DNA template and nuclease-free water (Sigma, St. Louis, USA). We included both positive controls (DNA from *L. major* - Friedlin str. and *L. donovani* (NLB065) and negative control (nuclease-free water) in each PCR reaction. We performed all the reactions in a SimpliAmp thermal cycler (Applied Biosystems, Loughborough, UK). The cycling conditions included an initial denaturation for 2 minutes at 98°C, followed by 35 cycles of denaturation for 20 seconds at 95°C, annealing for 30 seconds at 53°C, an extension for 30 seconds at 72°C and a final extension at 72°C for 5 minutes. The PCR products were run in 1.5% agarose gel stained with 1x ethidium bromide (Thermo Scientific, USA) and visualised in a GenoPlex gel documentation and analysis system (VWR, USA). The amplicons from positive samples were purified as aforementioned and submitted for sequencing at Macrogen (The Netherlands) using both the forward and reverse primers.

### Sandfly, *Leishmania*, and *Trypanosoma* sequence and phylogenetic analyses

The chromatograms of all sequences obtained were trimmed and edited using Geneious Prime software (v2020.0) to obtain consensus sequences for each sample. The consensus sequences were aligned using MUSCLE [31] with homologous sequences identified by sequence similarity searches in GenBank using the Basic Local Alignment Search Tool (BLAST) [32]. We constructed maximum-likelihood phylogenies of sandfly CO1, *Leishmania* ITS1, and *Trypanosoma* ITS1 sequence alignments using PhyML version 3.0 [33] employing the general-time-reversible (GTR) sequence evolution model. Tree-topologies were estimated over 1000 bootstrap replicates.

### Identification of vertebrate hosts represented in sandfly bloodmeals

We determined the bloodmeal sources of engorged sandflies by real-time PCR amplification of the vertebrate mitochondrial *cytochrome b (cyt-b)* gene followed by high-resolution melting (HRM) analysis as previously described [34]. All reactions were performed in a Rotor-Gene Q real-time PCR thermocycler (QIAGEN, Hannover, Germany) using DNA extracted from the blood of known vertebrate samples as positive controls, and nuclease-free water as the negative control. The positive controls included blood from livestock, rodents, and small mammals commonly found around the homestead such as rabbits and rock hyraxes. Livestock included goats, sheep, cow, and camels, while rodents included rats and mice. We also included human DNA extracted from blood obtained from a volunteer in *icipe*. Swiss mouse, rabbit, and rat blood were sourced from the *icipe*’*s* Animal Rearing Unit, while blood samples from the other animals were obtained from a slaughterhouse. We identified the vertebrate hosts represented in sandfly bloodmeals by comparing the melting profiles of the samples to those of the positive controls. Samples with melting profiles that did not match those of the positive controls were purified as described above and submitted for sequencing using the forward primer. The *cyt-b* chromatograms were edited in Geneious Prime software (v2020.0) and queried against the GenBank database using the NCBI’s BLASTn. The selected match was the top hit vertebrate species with the lowest e-value and homology cut-off values of 90%-100% as the most likely sandfly hosts.

### Statistical analyses

We performed statistical analyses using SPSS (v26) software. Descriptive statistics were used to determine the distribution pattern and frequency of each sandfly species per village. The species abundance was determined as the quantitative counts per village. The Kruskal-Wallis test was used to analyse differences in species distribution between villages.

## Results

### Sandfly species identification and distribution

We dissected a total of 526 sandflies, which were identified to the species level based on morphological keys. These consisted of one *Phlebotomus* (1.52%) and seven *Sergentomyia* species (98.48%) (Table 1). *Phlebotomus orientalis* was the only *Phlebotomus* species sampled during the study with the highest number being recorded in Matt Arban village (50%). The *Sergentomyia* species sampled in all the villages consisted of *Sergentomyia africanus* (0.19%), *S. antennatus* (0.38%), *S. bedfordi* (0.19%), *S. clydei* (10.46%), *S. inermis* (6.08%), *S. schwetzi* (2.47%), and *S. squamipleuris* (78.71%). *Sergentomyia squamipleuris* was the most abundant species in the study area, occurring in all the study villages. Sandfly distributions differed significantly (p < 0.001) across the sampled villages, with Kambi Juu village having the highest sandfly density. Most of the blood-fed sandflies were *Sergentomyia squamipleuris* (90%, n = 36), while the rest were *Phlebotomus orientalis* (n = 4).

**Table 1.** Sandfly abundance and density across the five sampling villages in Merti sub-County.

| Sandfly species | Trapping villages |  |  |  |  |  |  |  |  |  | Abundance |
| --- | --- | --- | --- | --- | --- | --- | --- | --- | --- | --- | --- |
|  | Kambi Juu |  | Barsa |  | Malkagala |  | Matt Arban |  | Korbesa |  |  |
|  | N | D | N | D | N | D | N | D | N | D |  |
| *Ph. orientalis* | 1 | 0.08 | 2 | 0.06 | 1 | 0.04 | 4 | 0.33 | 0 | 0.00 | 8 (1.52) |
| *S. africanus* | 0 | 0.00 | 0 | 0.00 | 0 | 0.00 | 1 | 0.08 | 0 | 0.00 | 1 (0.19) |
| *S. antennatus* | 0 | 0.00 | 1 | 0.03 | 1 | 0.04 | 0 | 0.00 | 0 | 0.00 | 2 (0.38) |
| *S. bedfordi* | 0 | 0.00 | 1 | 0.03 | 0 | 0.00 | 0 | 0.00 | 0 | 0.00 | 1 (0.19) |
| *S. clydei* | 7 | 0.58 | 42 | 1.16 | 6 | 0.25 | 0 | 0.00 | 0 | 0.00 | 55 (10.46) |
| *S. inermis* | 15 | 1.25 | 4 | 0.11 | 4 | 0.17 | 0 | 0.00 | 9 | 0.75 | 32 (6.08) |
| *S. schwetzi* | 12 | 1.00 | 1 | 0.03 | 0 | 0.00 | 0 | 0.00 | 0 | 0.00 | 13 (2.47) |
| *S. squamipleuris* | 90 | 7.50 | 50 | 1.39 | 159 | 6.63 | 104 | 8.67 | 11 | 0.91 | 414 (78.71) |
| Total | 125 | 10.41 | 101 | 2.81 | 171 | 7.13 | 109 | 9.08 | 20 | 1.66 | 526 (100) |

N: Sandfly counts; D: Sandfly density (counts/trap/night)

### Molecular identification of sandfly species

We amplified the *COI* region of 57 morphologically identified sandflies consisting of *Ph. orientalis, S. clydei*, and *S. squamipleuris* specimens. A BLAST search of the NCBI’s database using the *COI* gene sequences from the infected and a few uninfected *S. squamipleuris* sandflies (GenBank accessions: MT594454-MT594459) showed a high percentage identity (96%-99.8%) with those of *Sergentomyia* species in GenBank. Similarly, the *COI* sequences from *Ph. orientalis* (GenBank accessions: MT597050-MT597055) indicted a 97%-99.7% identity with other *Ph. orientalis* sequences in GenBank, thus confirming the morphological species identifications. Phylogenetic analysis of the *COI* sequences revealed genus-specific clusters with all the *Ph. orientalis*, and *S. squamipleuris* sequences separated into different branches (**Fig. 2**). The *S. squamipleuris* specimens from this study clustered together with other *S. squamipleuris* specimens collected from China (GenBank accessions: MF966747; MF966746). However, they segregated into different sub-branches which reflects the genetic isolation due to geography.

**Fig. 1.**
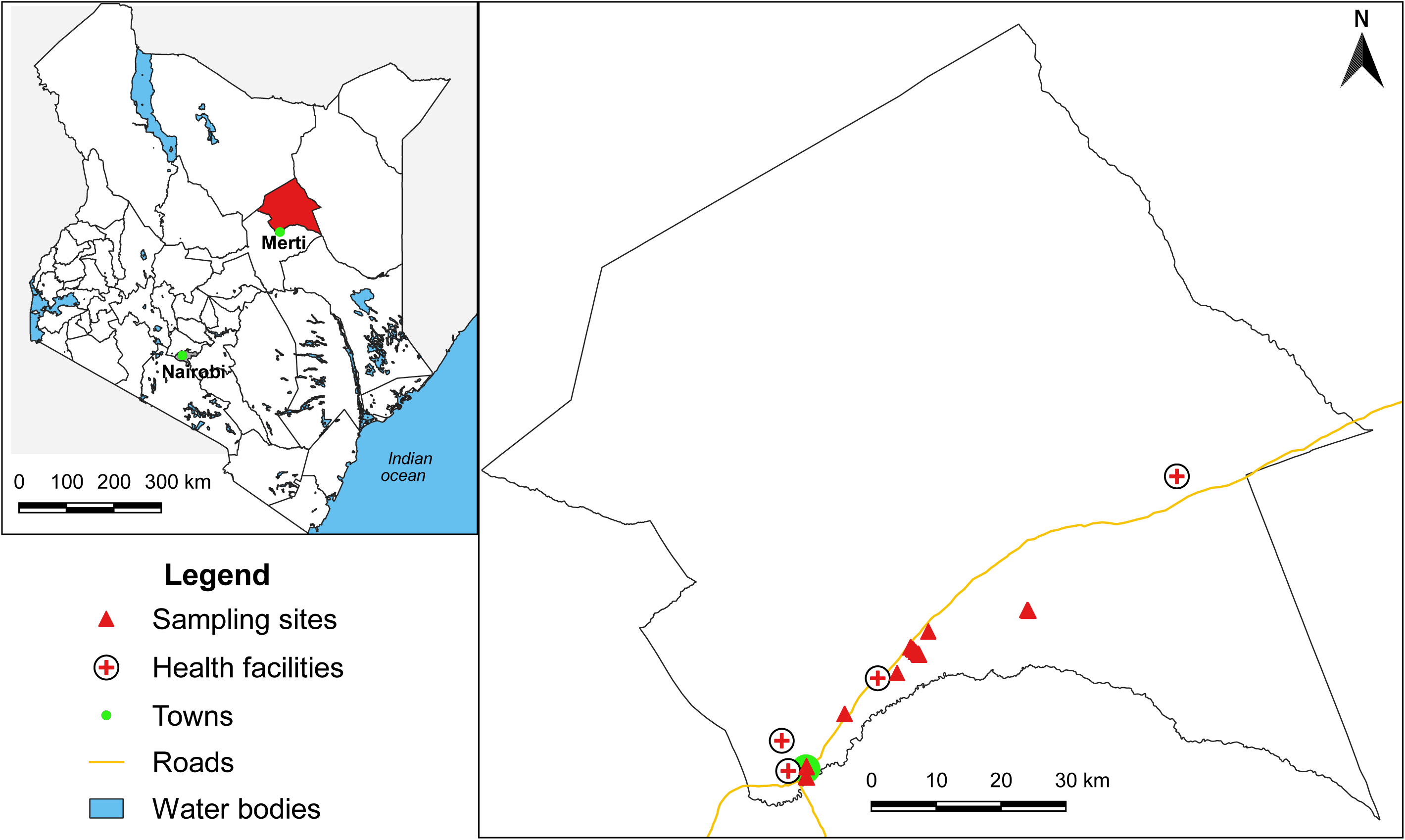
Left: Map of Kenya showing the geographical location of Merti sub-County. Right: Map of Merti sub-County showing the sampling sites. The maps were generated using QGIS version 3.0 [22].

**Fig. 2.**
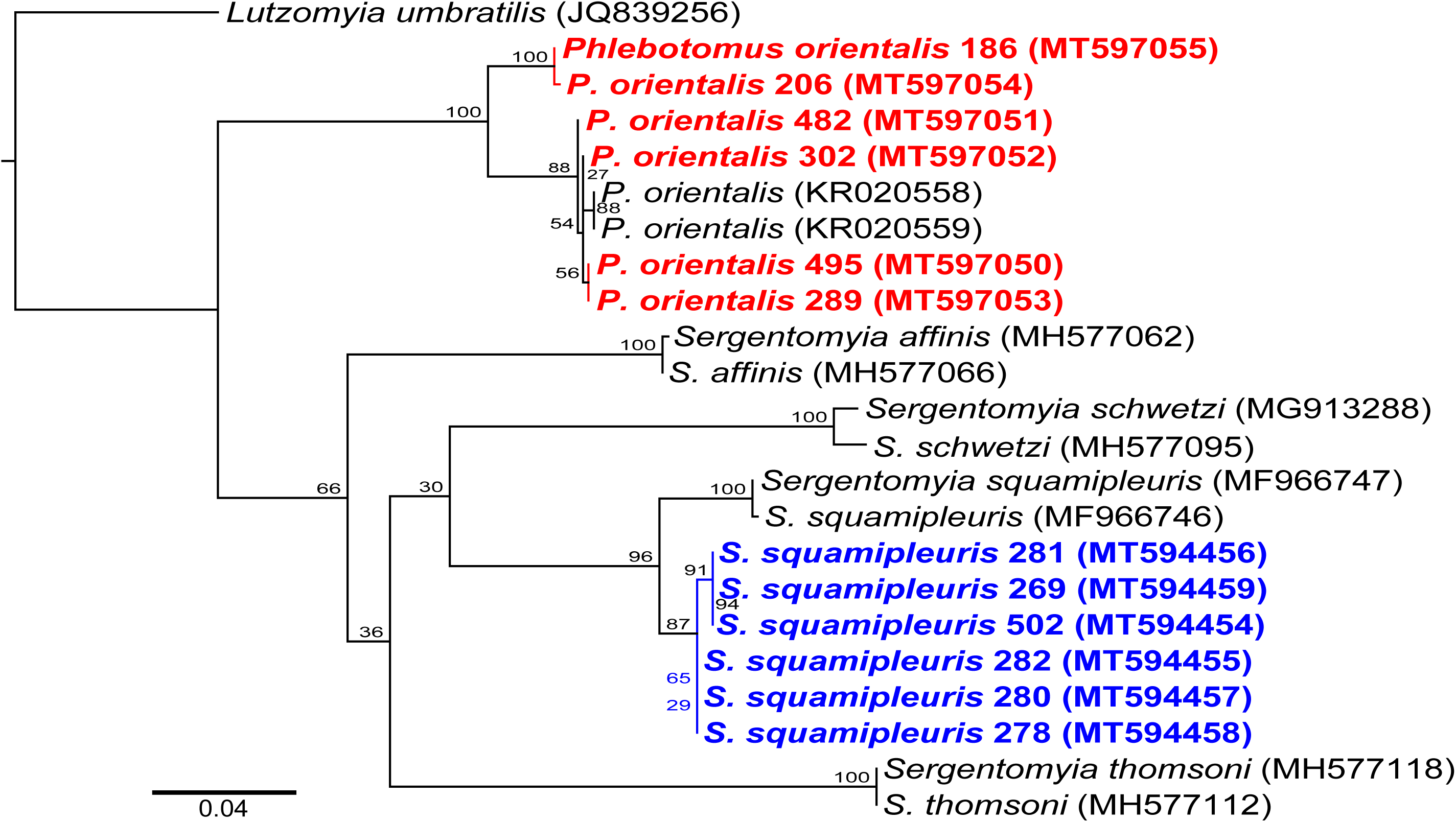
Maximum likelihood phylogenetic tree of sandfly COI gene sequences. The phylogeny was created from 700-bp fragments using New World sandfly species, *Lutzomyia umbratilis* (JQ839256), as an outgroup for rooting. The *Ph. orientalis* and *S. squamipleuris* sequences identified in this study are highlighted in bold. Bootstrap percentages at the major nodes are of agreement among 1000 replicates. The branch-length scale bar represents the number of nucleotide substitutions per site.

### Detection of *Leishmania* and *Trypanosoma* spp. in sandflies

We did not observe *Leishmania* promastigotes in the midguts of the dissected female sandflies by microscopy. To improve the sensitivity of *Leishmania* detection in the samples, we tested a total of 57 sandflies (Blood-fed: n = 40; unfed: 17) for the presence of *Leishmania* parasite’s DNA by PCR analysis of the *Leishmania* ITS1 region followed by sequencing. The samples consisted of *S. squamipleuris* (Blood-fed: 36; unfed: 13), *S. clydei* (unfed: 1) and *Ph. orientalis* (blood-fed: 4; unfed: 3). The *Leishmania* ITS1-PCR revealed four positive samples with bands of approximately 345-bp in one *S. squamipleuris* and approximately 320-bp in two *S. squamipleuris* specimens. Strikingly, we observed a band of approximately 570 bp in one *S. squamipleuris* specimen (**S1_Fig**). We did not detect *Leishmania* DNA in all *Ph. orientalis* and *S. clydei* samples analysed. More so, all the positive samples were collected from Malkagala village.

A BLAST search of the NCBI’s database using the *ITS1* sequence (GenBank accession: MT548852) from one of the positive *S. squamipleuris* samples revealed a high percentage identity (98.8%) with *L. major* sequences in GenBank. Similarly, sequences from two of the positive *S. squamipleuris* samples (GenBank accessions: MT548853-MT548854) showed a 96.9-100% identity with *L. donovani* sequences in GenBank. In the phylogenetic tree, the *L. major* and *L. donovani* ITS1 sequences from this study segregated into the *Leishmania major* and *Leishmania donovani* clusters with bootstrap support values of 92% and 99%, respectively (**Fig. 3**). These clusters contained *Leishmania* spp. that have been isolated from patients in different regions and whose sequences are available in GenBank.

**Fig. 3.**
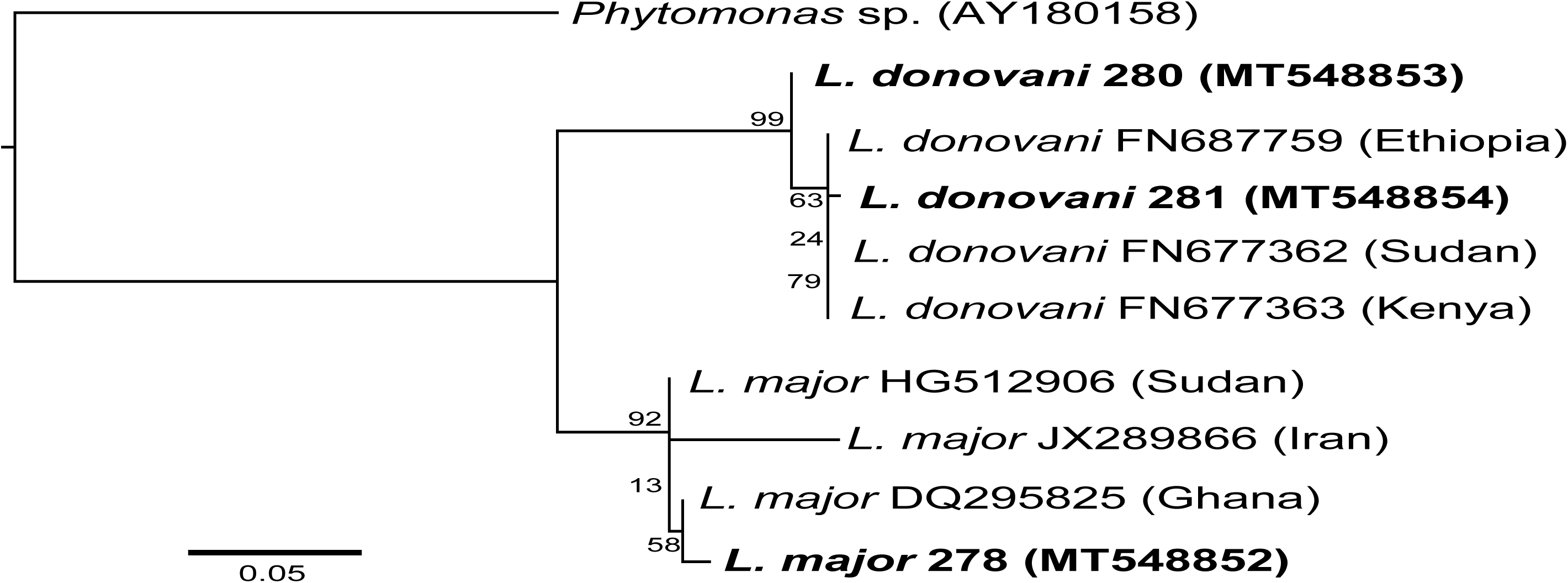
Maximum likelihood phylogenetic tree of *Leishmania* ITS1 gene sequences. The phylogeny was created from 345-bp and 320-bp fragments, for *L. major* and *L. donovani*, respectively, using *Phytomonas* sp. (AY180158) as an outgroup for rooting. The *L. major* and *L. donovani* sequences identified in this study are highlighted in bold. Bootstrap percentages at the major nodes are of agreement among 1000 replicates. The branch-length scale bar represents the number of nucleotide substitutions per site.

Unexpectedly, the ITS1 sequence of the 570-bp amplicon (GenBank accession: MT548851) did not match those of *Leishmania* spp., but rather showed high homology with *Trypanosoma* spp., suggesting that it belonged to the *Trypanosoma* genus. A BLAST search analysis of the sequence revealed a 91.6% identity with the ITS1 sequence of *Trypanosoma rangeli* registered in the GenBank. This sequence was placed into a monophyletic cluster with other trypanosomes isolated from humans and other animals and was unrelated to those isolated from reptiles (e.g. *T. grayi*). We found the sequence to be closely related to that of *T. rangeli* isolated from humans than the other *Trypanosoma* species (**Fig. 4**), which further suggests the subgenera it belongs to. Although this finding was further corroborated by patristic distance matrix analysis (**S2_Fig**.), the *Trypanosoma* DNA detected within *Sergentomyia squamipleuris* segregated into a different monophyletic sub-branch, suggesting that it was novel or genetically uncharacterised.

**Fig. 4.**
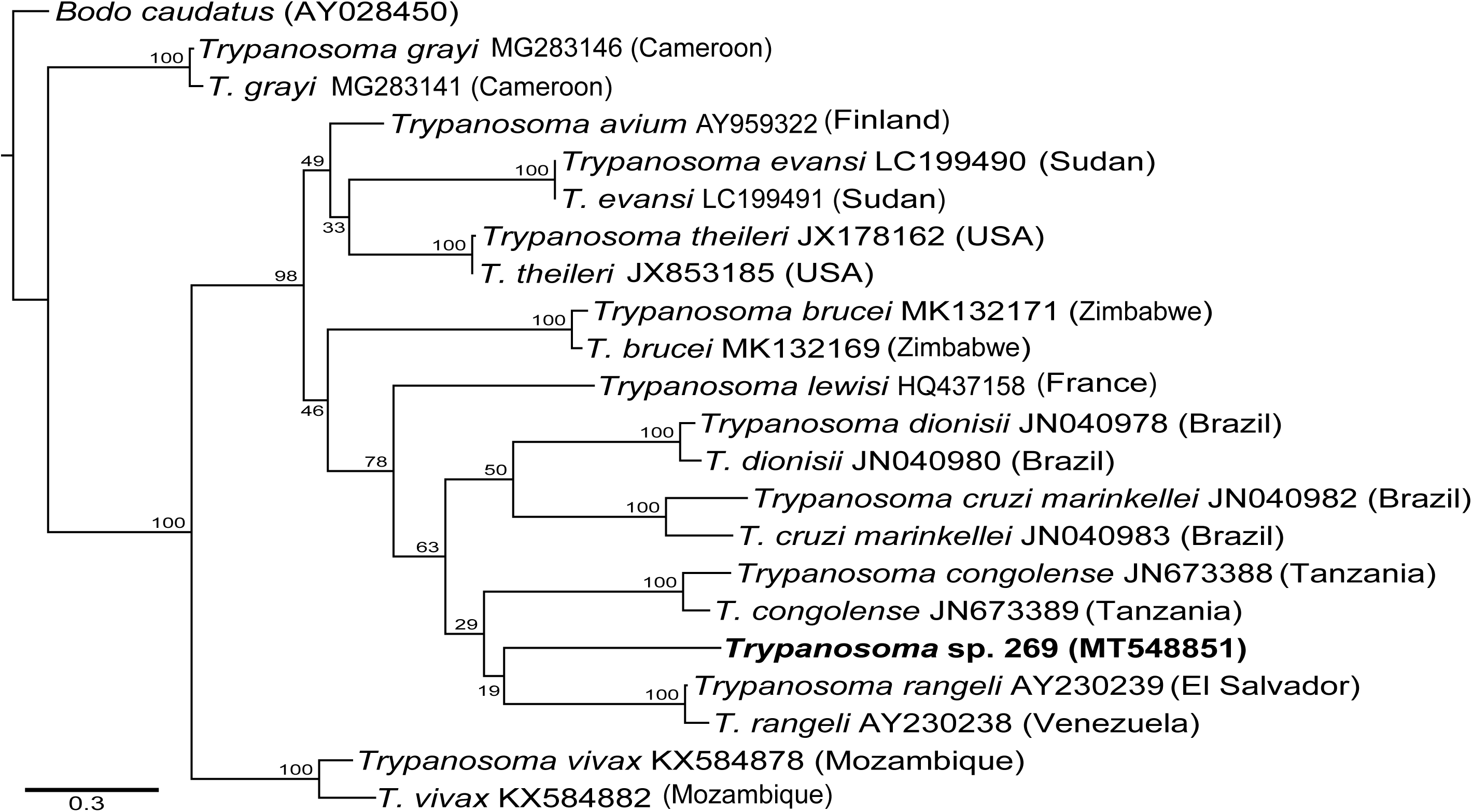
Maximum likelihood phylogenetic tree of *Trypanosoma* species ITS1 gene sequences. The phylogeny was created from 570-bp fragments using *Bodo caudatus* (AY028450) as an outgroup for rooting. The *Trypanosoma* sequence identified in this study is highlighted in bold. Bootstrap percentages at the major nodes are of agreement among 1000 replicates. The branch-length scale bar represents the number of substitutions per site.

### Sandfly bloodmeal source determination

We determined the bloodmeal sources of all the engorged sandflies consisting of *S. squamipleuris* (n = 36) and four *Ph. orientalis* sandflies. The *cyt-b* HRM profiles revealed various vertebrate hosts in sandfly bloodmeals, including humans (*Homo sapiens*), rock hyraxes (*Procavia capensis*), and rodents (**Table 2**). Humans were the main bloodmeal sources detected exclusively in 66.7% (n = 24) and 50% (n = 2) of the engorged *S. squamipleuris* and *Ph. orientalis*, respectively. All the sandflies that were infected with *Leishmania major* and *Leishmania donovani* had taken a bloodmeal exclusively from humans. The presence of mixed feeding was identified based on HRM profiles with multiple peaks compared to the reference controls. We identified mixed feeding in three samples: two *Phlebotomus orientalis* and one *Sergentomyia squamipleuris*. One of the *Phlebotomus orientalis* specimens had taken a blood meal from humans and rock hyraxes while the other had multiple bloodmeals amongst humans, rock hyraxes and mice. In contrast, the *Sergentomyia squamipleuris* specimen had bloodmeals from humans and mice. We did not characterise bloodmeal sources of 11 samples including one sample that we found to be infected with *Trypanosoma* spp. as the *cyt-b* amplification failed in these samples. The representative *cyt-b* sequences from this study have been deposited in the GenBank nucleotide database under the accession numbers: MT568790-MT568795.

**Table 2:**
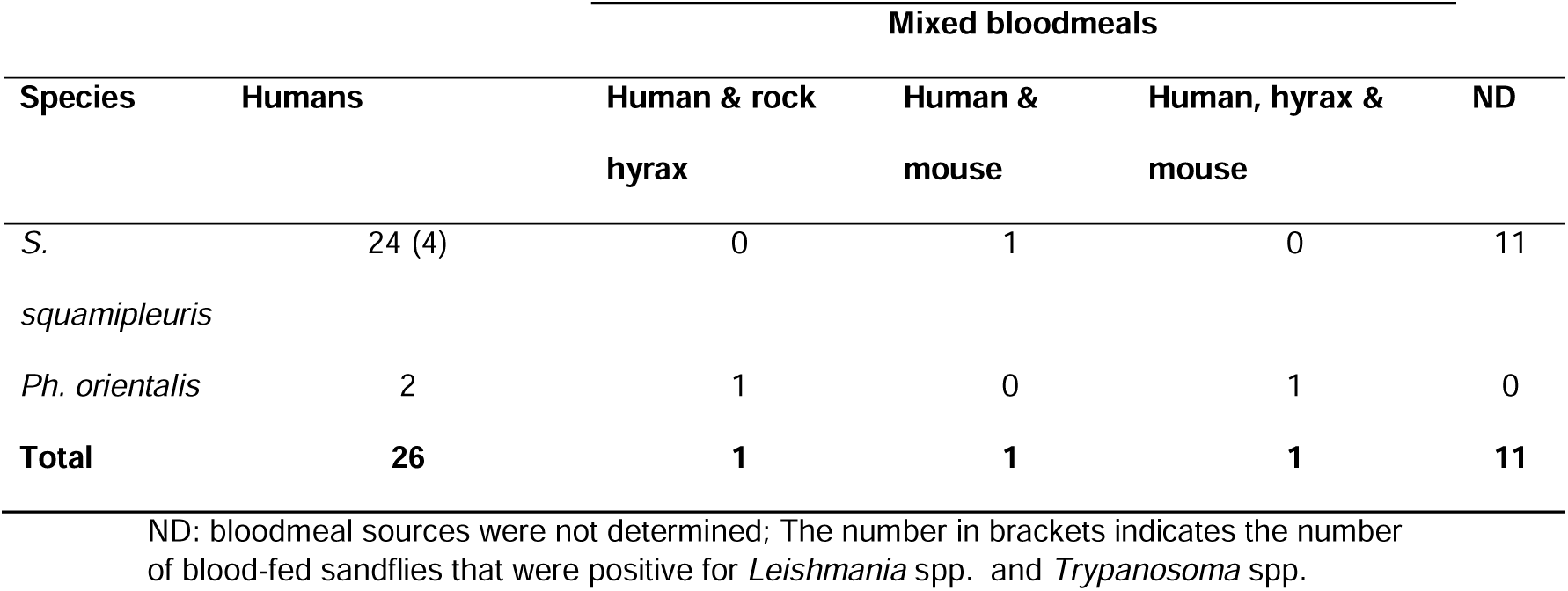
sandfly bloodmeal sources in Merti sub-County determined by real-time PCR of the vertebrate *cyt-b* gene followed by HRM analysis.

## Discussion

The main goal of this study was to examine sandfly species with the potential to transmit *Leishmania* parasites in Merti sub-County and determine their abundance, distribution, and bloodmeal sources. We detected *L. major, L. donovani*, and *Trypanosoma* spp., for the first time, in *Sergentomyia squamipleuris* from a VL-endemic area in Kenya. These findings reveal urgent questions to be addressed regarding the dogma that human leishmaniases in the Old World are exclusively transmitted by sandflies of the *Phlebotomus* genus [1,35]

We identified eight sandfly species from all the study villages. These consisted of one *Phlebotomus* and seven *Sergentomyia* species. *Phlebotomus orientalis* was the only *Phlebotomus* species sampled in this study and represented one out of the five *Phlebotomus* subgenera described in Kenya [25]. *Phlebotomus orientalis* is implicated as the secondary vector of *L. donovani* in Kenya [10]. However, it was found to occur in relatively low abundance in all the sampling villages. The low numbers of *Ph. orientalis* in this study corroborated previous reports in Kenya that suggest that the species do not occur in large numbers in VL endemic areas [10]. The low abundance of *Ph. orientalis* could be attributed to very few cracked vertisols in the area during the sampling period. Since we conducted the sampling after the long rains season, the soil had not dried enough to form deep cracks which act as breeding sites for the sandfly species [28]. Sandflies of the *Sergentomyia* subgenus were the predominant species, with *S. squamipleuris* being the most abundant, accounting for 78.71% and present in all the study villages, followed by *S. clydei* (10.46%). We found the distribution of sandfly species across the sampling villages to be significantly different with the highest sandfly density being recorded in Kambi Juu, while Korbesa had the lowest sandfly density.

Phylogenetic analysis of the sandfly *COI* sequences separated *Ph. orientalis* and *S. squamipleuris* into genus-specific clusters, confirming the morphological species identifications. The high prevalence of *Leishmania* spp. and the detection of *Trypanosoma* spp. in *S. squamipleuris* sandflies suggest the possible involvement of the sandfly in the transmission of *Leishmania* spp. and *Trypanosoma* spp. in Merti.

Demonstrating *Leishmania* infections in sandflies is a prerequisite for vector incrimination. The vectorial role of a sandfly species is epidemiologically suspected if the species is predominant in a leishmaniasis focus and exhibits an anthropophilic and anthropophagic behaviour [15]. This hypothesis is supported if the same species is found infected with live transmissible promastigotes like those isolated from humans and other vertebrate reservoirs. In this study, we did not observe *Leishmania* promastigotes in the midguts of all the dissected female sandflies by microscopy. Although this method is used as the gold standard for demonstrating *Leishmania* infections in sandflies [36], it has low sensitivity, which reduces with a decrease in parasite burden in the flies. The low sensitivity could explain why we did not observe live infections in the infected flies. Using the ITS1-PCR followed by sequencing, we detected, for the first time in Kenya, the *Leishmania* spp. and *Trypanosoma* spp. DNA in four samples of *S. squamipleuris. Leishmania major* and *Trypanosoma* species were detected in one specimen each, while two of the samples were found to harbour *L. donovani* parasites.

Comparison of *Leishmania* spp. identified in sandflies with those isolated from humans is crucial in predicting the risks of disease transmissions in endemic areas. Phylogenetic analysis of the ITS1 sequences of the *Leishmania* spp. identified in this study revealed that the sequences were closely related to other *Leishmania* species isolated from humans. The *L. major* sequence segregated into the *L. major* cluster with bootstrap support values of 100% and was closely related to *L. major* sequence isolated from a CL patient in Ghana [37]. Similarly, we found the *L. donovani* sequence to be closely related to other *L. donovani* sequences isolated from humans. However, the two sequences were separated into monophyletic clusters which may suggest an incomplete lineage sorting. The detection of *L. major* and *L. donovani*, similar to those infecting humans, in *S. squamipleuris* sandflies from Merti focus is a novel finding and corroborates other studies in Kenya [19] and elsewhere [13,15,16,18], in which sandflies of this genus were found to be naturally infected with *Leishmania* spp. pathogenic to humans.

To date, to our knowledge, there is no published report on the role of *S. squamipleuris* as a vector of *Leishmania* or *Trypanosoma* spp. in Kenya. Our results, in addition to the previous reports, further suggest that *S. squamipleuris* could be involved in the transmission cycle of *Leishmania* parasites in Merti, where there were low numbers of *Phlebotomus* species and none of these was found to be infected with *Leishmania* parasites. Although Sadlova and colleagues [12] reported that *Leishmania* spp. pathogenic to humans cannot develop in *S. schwetzi*, we found *L. major* and *L. donovani* infections in *S. squamipleuris*. Further studies are needed to establish the competence of *S. squamipleuris* to transmit *L. major* and *L. donovani*. Furthermore, the detection of *Trypanosoma* spp. in *S. squamipleuris* in this study supports a previous study in which *Trypanosoma* spp. was also identified in *S. africana africana* in Ghana [18]. Further investigations are needed to determine whether *Sergentomyia* sandflies can support the cyclic development of *Trypanosoma* spp. that are pathogenic to humans and other animals.

Analysis of the sandfly bloodmeals sources revealed humans as the main sandfly bloodmeal hosts in Merti sub-County. The significant preference for humans (65%) further suggests a high risk of disease transmissions in the study site and could explain the recurrent outbreaks and the rising number of VL cases in the area. Most of the blood-fed sandflies were *S. squamipleuris* (90%), the majority of which had taken bloodmeals exclusively from humans. Only two of the *S. squamipleuris* specimens had mixed bloodmeals in their gut. These results aligned well with other studies in Sudan where sandflies of the *Sergentomyia* genus were reported as a common human-biting species [13]. Of the 24 *S. squamipleuris* sandflies that fed on humans, two were infected with *L. donovani*, one with *L. major* and one with *Trypanosoma* spp. These findings challenge the dogma that human leishmaniases in the Old World are exclusively transmitted by sandflies of the *Phlebotomus* genus [1,35]. Although, we identified human bloodmeals in two *Ph. orientalis* samples and mixed feeding in two additional samples of the same species, none of these samples were found to be infected with *Leishmania* parasites. Our results illustrate the potential role of *S. squamipleuris* in the transmission of *Leishmania* spp. in Kenya, providing useful information to guide in addressing the epidemiology of the disease including control.

Other vertebrate hosts represented in sandfly bloodmeals included rock hyrax (*Procavia capensis*) and mouse (*Mus musculus*). Since both *Ph. orientalis* and *S. squamipleuris* do not feed exclusively on humans, there is a possibility of zoonotic transmission of leishmaniasis and other pathogens in this VL focus. Characterisation of bloodmeal sources failed in 11 samples. This could be due to the degradation of the *cyt-b* target, highlighting the importance of using more than one genetic marker for the analyses of host preferences.

Detection of both *L. donovani* and *L. major* in *S. squamipleuris* that mainly fed on humans further suggests that this species may be a potential permissive vector of the *Leishmania* species. In most permissive vectors of *Leishmania* spp. such as *Lu. longipalpis* and *Ph. arabicus*, the attachment of *Leishmania* parasites to the midgut is independent of the midgut lipophosphoglycan (LPG). Whether such mechanisms also apply to the sandflies of the *Sergentomyia* species in which more than one *Leishmania* spp. pathogenic to humans have been identified remains to be determined. Moreover, further studies are needed to determine the role of *S. squamipleuris* in the transmission cycle of *L. donovani* and *L. major* parasites by determining the developmental stages of the parasites in individual field-collected sandflies and transmission experiments or through evaluating *Leishmania* developmental stage-specific gene expression.

## Conclusions

We demonstrate, for the first time, the detection of *L. major, L. donovani*, and *Trypanosoma* spp. in *S. squamipleuris* sandflies from a VL focus in Merti sub-County, eastern Kenya. This sandfly species was the most abundant in the study area and exhibited anthropophilic and anthropophagic behaviour, which suggests its potential involvement in *Leishmania* transmission in the sub-County. Identification of both *L. major* and *L. donovani* in *S. squamipleuris* further suggests that it may be a potential permissive vector of both parasites, a finding that needs further investigations. Identification of *Trypanosoma* spp. in *S. squamipleuris* may indicate mechanical transmission, whose efficiency should be investigated. Analysis of sandfly bloodmeal sources revealed the possibility of zoonotic transmission of leishmaniasis and possibly other pathogens in Merti sub-County. Further studies are needed to incriminate the vector and establish vector competence and reservoir hosts of *Leishmania* parasites in this area.

## Supporting information

**S1_Fig. Detection of *Leishmania* and *Trypanosoma* spp. DNA in sand flies by ITS1-PCR**. M: 100 bp ladder; 1-4: *Sergentomyia squamipleuris* sandfly samples; 5 and 6: *Leishmania donovani* (NLB065) and *Leishmania major* (Friedlin strain) positive controls; NTC: negative control.

**S2_Fig**. Summary matrix of pair-wise patristic distances between the ITS1 sequences of *Trypanosoma* spp. identified in this study (red borders) and those registered in the GenBank. The patristic distances between a pair of sequences are represented in the form of a heatmap and their values indicated. The genetic distances were calculated in geneious prime (v20.0) following a maximum likelihood phylogenetic analysis implemented in PHYML.

## Abbreviations

CDC: Centres for Disease Control and Prevention
COI: Cytochrome c oxidase subunit 1
*cyt-b*: cytochrome b
DNA: deoxyribonucleic acid
GTR: General-time-reversible
HRM: high-resolution melt
*icipe*: International Centre of Insect Physiology and Ecology
ITS1: internal transcribed spacer 1
KEMRI: Kenya Medical Research Institute
NTDs: Neglected Tropical Diseases
PCR: Polymerase Chain Reaction
VL: Visceral Leishmaniasis
ZCL: zoonotic cutaneous leishmaniasis.

## Acknowledgements

We thank administrative officers, the community health workers in Merti sub-County and the Ministry of Health for support in the implementation of fieldwork and logistics. We acknowledge the Director-General, KEMRI and the Deputy Director CBRD for providing laboratory workspace, personnel, and logistical support in the implementation of the project. We also acknowledge the KEMRI grantsmanship office, who provided funding for the project. We appreciate the support from the *Leishmania* section, KEMRI, including the late Dr Christopher Anjili, Dr Josyline Kaburi, Bernard Ongondo, Reuben Ruttoh, David Siele, Joash Mandere and Newton Mwathi. We acknowledge Karen Wambui, Faith Kyengo and Gerard Ronoh of *icipe* for technical assistance and help with the logistics.

## Ethics approval and consent to participate

The study was approved by the KEMRI Scientific Ethical Review Committee (SERU), KEMRI/SERU/CBRD/164/3447 and informed oral consent was obtained from household owners before sandfly collection.

## Availability of Data and materials

All the data generated or analysed during this study are included in this published article. The datasets used and/or analysed during the current study are available from the corresponding author on reasonable request. Selected sandfly, parasite and bloodmeal sequences generated and/or analysed during the current study are available in the GenBank repository: *COI* (MT597050-MT597055, MT594454-MT594459); ITS1 (MT548851-MT548854); *cyt-b* (MT568790-MT568795)

## Funding

This study was supported by the KEMRI IRG grant (KEMR1/1RG/165/6 awarded to JMM), Kenya National Research Fund (awarded to DKM & DMM), Wellcome Trust grants (080883/E/06/A awarded to DMM and 205304/Z/16/Z awarded to BOO), and AntiVec grant (AV/PP0018/1 awarded to JV & DMM). Additionally, we gratefully acknowledge the financial support for this research by the following organizations and agencies: UK’s Department for International Development (DFID); the Swedish International Development Cooperation Agency (Sida); the Swiss Agency for Development and Cooperation (SDC); the government of the Republic of Kenya, and the government of the Republic of Ethiopia. The funders had no role in study design, data collection and analysis, decision to design, data collection and analysis, decision to publish, or preparation of the manuscript.

## Author contributions

Conceptualization: JMM, JV, DKM, DMM; Data curation: BOO, JMM, SK, HNM, AC, JMI, DMM; Formal analysis: BOO, JMM, SK, HNM, JMI, AC, PMN, JV, DKM, DMM; Funding acquisition: JMM, JV, DKM, DMM; Investigation: BOO, JMM, JV, DKM, DMM; Methodology: BOO, JMM, SK, HN, JMI, AC, PMN, JV, DKM, DMM; Project administration: JV, DKM, DMM. Supervision: JV, DKM, DMM. Validation: BOO, JMM, SK, HNM, JMI, AC, PMN, JV, DKM, DMM. Visualization: BOO, JMM, SK, HNM, JMI, AC, PMN, JV, DKM, DMM; Manuscript drafting: BOO, JMM; Review & editing: BOO, JMI, JMM, PMN, JV, DKM, DMM.

## Competing interests

The authors declare that they have no competing interest

## Consent for publication

Not applicable

## Notes

### Competing Interest Statement

The authors have declared no competing interest.

