## Supplementary figures and images for "Molecular detection of *Leishmania donovani, Leishmania major*, and *Trypanosoma* spp. in *Sergentomyia squamipleuris* sandflies from a visceral leishmaniasis focus in Merti sub-County, eastern Kenya"

### Supplementary figure 1

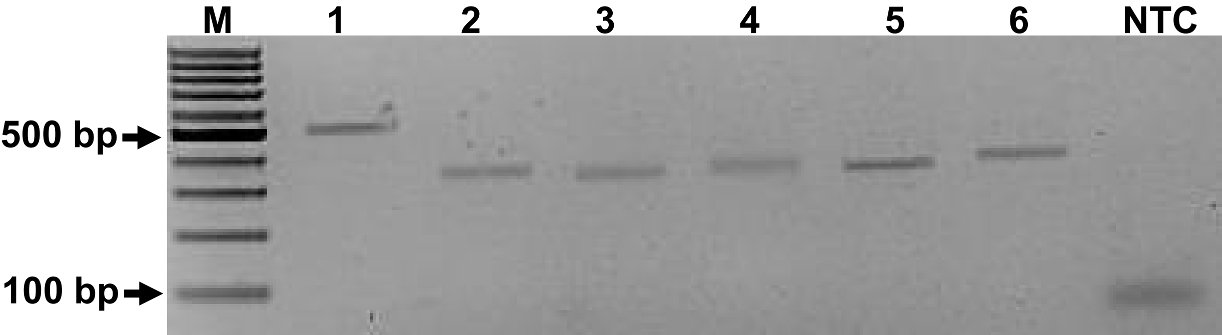

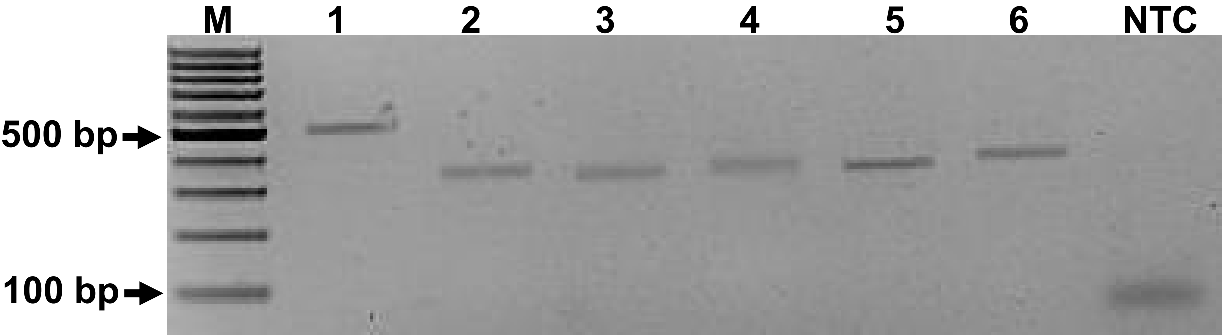
