## Supplementary figure 2 for "Molecular detection of *Leishmania donovani, Leishmania major*, and *Trypanosoma* spp. in *Sergentomyia squamipleuris* sandflies from a visceral leishmaniasis focus in Merti sub-County, eastern Kenya"

| | *Trypanosoma* spp. 269 (MT548851) | *T. dionisii* (JN040980.1) | *T. dionisii* (JN040978.1) | *T. cruzi* (JN040983.1) | *T. cruzi* (JN040982.1) | *T. lewisi* (HQ437158.1) | *T. avium* (AY959322.1) | *T. theileri* (JX178162.1) | *T. theileri* (JX853185.1) | *T. vivax* (KX584882.1) | *T. vivax* (KX584878.1) | *T. evansi* (LC199491.1) | *T. evansi* (LC199490.1) | *Bodo caudatus* (AY028450.1) | *T. grayi* (MG283141.1) | *T. grayi* (MG283146.1) | *T. rangeli* (AY230239.1) | *T. rangeli* (AY230238.1) | *T. congolense* (JN673388.1) | *T. congolense* (JN673389.1) | *T. brucei* (MK132171.1) | *T. brucei* (MK132169.1) |
|---|---|---|---|---|---|---|---|---|---|---|---|---|---|---|---|---|---|---|---|---|---|---|
| *Trypanosoma* spp. 269 (MT548851) | | 1.2 | 1.2 | 1.3 | 1.3 | 1.2 | 1.1 | 1.3 | 1.3 | 1.6 | 1.5 | 1.5 | 1.5 | 1.7 | 1.9 | 1.8 | 0.9 | 0.9 | 1.1 | 1.0 | 1.5 | 1.4 |
| *T. dionisii* (JN040980.1) | 1.2 | | 0.1 | 0.9 | 0.9 | 1.1 | 1.1 | 1.3 | 1.3 | 1.6 | 1.5 | 1.5 | 1.5 | 1.6 | 1.8 | 1.8 | 1.1 | 1.2 | 1.2 | 1.2 | 1.4 | 1.4 |
| *T. dionisii* (JN040978.1) | 1.2 | 0.1 | | 0.9 | 0.9 | 1.1 | 1.1 | 1.3 | 1.3 | 1.5 | 1.5 | 1.4 | 1.4 | 1.6 | 1.8 | 1.8 | 1.1 | 1.1 | 1.2 | 1.2 | 1.4 | 1.4 |
| *T. cruzi* (JN040983.1) | 1.3 | 0.9 | 0.9 | | 0.3 | 1.2 | 1.1 | 1.3 | 1.3 | 1.6 | 1.6 | 1.5 | 1.5 | 1.7 | 1.9 | 1.9 | 1.2 | 1.2 | 1.3 | 1.2 | 1.5 | 1.5 |
| *T. cruzi* (JN040982.1) | 1.3 | 0.9 | 0.9 | 0.3 | | 1.2 | 1.2 | 1.4 | 1.4 | 1.7 | 1.6 | 1.6 | 1.6 | 1.7 | 1.9 | 1.9 | 1.2 | 1.3 | 1.3 | 1.3 | 1.5 | 1.5 |
| *T. lewisi* (HQ437158.1) | 1.2 | 1.1 | 1.1 | 1.2 | 1.2 | | 0.8 | 1.0 | 1.0 | 1.3 | 1.3 | 1.2 | 1.2 | 1.4 | 1.6 | 1.5 | 1.1 | 1.2 | 1.2 | 1.2 | 1.2 | 1.2 |
| *T. avium* (AY959322.1) | 1.1 | 1.1 | 1.1 | 1.1 | 1.2 | 0.8 | | 0.4 | 0.4 | 0.8 | 0.8 | 0.6 | 0.6 | 0.9 | 1.1 | 1.1 | 1.0 | 1.1 | 1.1 | 1.1 | 0.8 | 0.8 |
| *T. theileri* (JX178162.1) | 1.3 | 1.3 | 1.3 | 1.3 | 1.4 | 1.0 | 0.4 | | 0.0 | 1.0 | 1.0 | 0.7 | 0.7 | 1.1 | 1.3 | 1.3 | 1.2 | 1.3 | 1.3 | 1.3 | 1.0 | 1.0 |
| *T. theileri* (JX853185.1) | 1.3 | 1.3 | 1.3 | 1.3 | 1.4 | 1.0 | 0.4 | 0.0 | | 1.0 | 1.0 | 0.7 | 0.7 | 1.1 | 1.3 | 1.3 | 1.2 | 1.3 | 1.3 | 1.3 | 1.0 | 1.0 |
| *T. vivax* (KX584882.1) | 1.6 | 1.6 | 1.5 | 1.6 | 1.7 | 1.3 | 0.8 | 1.0 | 1.0 | | 0.2 | 1.2 | 1.2 | 0.9 | 1.1 | 1.1 | 1.5 | 1.6 | 1.6 | 1.6 | 1.3 | 1.3 |
| *T. vivax* (KX584878.1) | 1.5 | 1.5 | 1.5 | 1.6 | 1.6 | 1.3 | 0.8 | 1.0 | 1.0 | 0.2 | | 1.2 | 1.2 | 0.8 | 1.0 | 1.0 | 1.5 | 1.5 | 1.6 | 1.5 | 1.2 | 1.2 |
| *T. evansi* (LC199491.1) | 1.5 | 1.5 | 1.4 | 1.5 | 1.6 | 1.2 | 0.6 | 0.7 | 0.7 | 1.2 | 1.2 | | 0.0 | 1.3 | 1.5 | 1.5 | 1.4 | 1.5 | 1.5 | 1.5 | 1.2 | 1.2 |
| *T. evansi* (LC199490.1) | 1.5 | 1.5 | 1.4 | 1.5 | 1.6 | 1.2 | 0.6 | 0.7 | 0.7 | 1.2 | 1.2 | 0.0 | | 1.3 | 1.5 | 1.5 | 1.4 | 1.5 | 1.5 | 1.5 | 1.2 | 1.2 |
| *Bodo caudatus* (AY028450.1) | 1.7 | 1.6 | 1.6 | 1.7 | 1.7 | 1.4 | 0.9 | 1.1 | 1.1 | 0.9 | 0.8 | 1.3 | 1.3 | | 0.5 | 0.5 | 1.6 | 1.6 | 1.7 | 1.6 | 1.4 | 1.4 |
| *T. grayi* (MG283141.1) | 1.9 | 1.8 | 1.8 | 1.9 | 1.9 | 1.6 | 1.1 | 1.3 | 1.3 | 1.1 | 1.0 | 1.5 | 1.5 | 0.5 | | 0.0 | 1.8 | 1.8 | 1.9 | 1.8 | 1.6 | 1.5 |
| *T. grayi* (MG283146.1) | 1.8 | 1.8 | 1.8 | 1.9 | 1.9 | 1.5 | 1.1 | 1.3 | 1.3 | 1.1 | 1.0 | 1.5 | 1.5 | 0.5 | 0.0 | | 1.8 | 1.8 | 1.9 | 1.8 | 1.5 | 1.5 |
| *T. rangeli* (AY230239.1) | 0.9 | 1.1 | 1.1 | 1.2 | 1.2 | 1.1 | 1.0 | 1.2 | 1.2 | 1.5 | 1.5 | 1.4 | 1.4 | 1.6 | 1.8 | 1.8 | | 0.1 | 1.0 | 1.0 | 1.4 | 1.4 |
| *T. rangeli* (AY230238.1) | 0.9 | 1.2 | 1.1 | 1.2 | 1.3 | 1.2 | 1.1 | 1.3 | 1.3 | 1.6 | 1.5 | 1.5 | 1.5 | 1.6 | 1.8 | 1.8 | 0.1 | | 1.1 | 1.0 | 1.4 | 1.4 |
| *T. congolense* (JN673388.1) | 1.1 | 1.2 | 1.2 | 1.3 | 1.3 | 1.2 | 1.1 | 1.3 | 1.3 | 1.6 | 1.6 | 1.5 | 1.5 | 1.7 | 1.9 | 1.9 | 1.0 | 1.1 | | 0.2 | 1.5 | 1.5 |
| *T. congolense* (JN673389.1) | 1.0 | 1.2 | 1.2 | 1.2 | 1.3 | 1.2 | 1.1 | 1.3 | 1.3 | 1.6 | 1.5 | 1.5 | 1.5 | 1.6 | 1.8 | 1.8 | 1.0 | 1.0 | 0.2 | | 1.4 | 1.4 |
| *T. brucei* (MK132171.1) | 1.5 | 1.4 | 1.4 | 1.2 | 1.5 | 1.2 | 0.8 | 1.0 | 1.0 | 1.3 | 1.2 | 1.2 | 1.2 | 1.4 | 1.6 | 1.5 | 1.4 | 1.4 | 1.5 | 1.4 | | 0.1 |
| *T. brucei* (MK132169.1) | 1.4 | 1.4 | 1.4 | 1.5 | 1.5 | 1.2 | 0.8 | 1.0 | 1.0 | 1.3 | 1.2 | 1.2 | 1.2 | 1.4 | 1.5 | 1.5 | 1.4 | 1.4 | 1.5 | 1.4 | 0.1 | |
